# SARS-CoV-2 Simulations Go Exascale to Capture Spike Opening and Reveal Cryptic Pockets Across the Proteome

**DOI:** 10.1101/2020.06.27.175430

**Authors:** Maxwell I. Zimmerman, Justin R. Porter, Michael D. Ward, Sukrit Singh, Neha Vithani, Artur Meller, Upasana L. Mallimadugula, Catherine E. Kuhn, Jonathan H. Borowsky, Rafal P. Wiewiora, Matthew F. D. Hurley, Aoife M Harbison, Carl A Fogarty, Joseph E. Coffland, Elisa Fadda, Vincent A. Voelz, John D. Chodera, Gregory R. Bowman

## Abstract

SARS-CoV-2 has intricate mechanisms for initiating infection, immune evasion/suppression, and replication, which depend on the structure and dynamics of its constituent proteins. Many protein structures have been solved, but far less is known about their relevant conformational changes. To address this challenge, over a million citizen scientists banded together through the Folding@home distributed computing project to create the first exascale computer and simulate an unprecedented 0.1 seconds of the viral proteome. Our simulations capture dramatic opening of the apo Spike complex, far beyond that seen experimentally, which explains and successfully predicts the existence of ‘cryptic’ epitopes. Different Spike homologues modulate the probabilities of open versus closed structures, balancing receptor binding and immune evasion. We also observe dramatic conformational changes across the proteome, which reveal over 50 ‘cryptic’ pockets that expand targeting options for the design of antivirals. All data and models are freely available online, providing a quantitative structural atlas.

## Introduction

Severe acute respiratory syndrome coronavirus 2 (SARS-CoV-2) is a novel coronavirus that poses an imminent threat to global human health and socioeconomic stability.^1^ With estimates of the basic reproduction number at ∼3-4 and a case fatality rate for coronavirus disease 2019 (COVID-19) ranging from ∼0.1-12% (high temporal variation), SARS-CoV-2/COVID-19 has spread quickly and currently endangers the global population.^2-6^ As of September 12^th^, 2020, there have been over 29 million confirmed cases and over 925,000 fatalities, globally. Quarantines and social distancing are effective at slowing the rate of transmission; however, they cause significant social and economic disruption. Taken together, it is crucial that we find immediate therapeutic interventions.

A structural understanding of the SARS-CoV-2 proteins could accelerate the discovery of new therapeutics by enabling the use of rational design.^7^ Towards this end, the structural biology community has made heroic efforts to rapidly build models of SARS-CoV-2 proteins and the complexes they form. However, it is well established that a protein’s function is dictated by the full range of conformations it can access; many of which remain hidden to experimental methods. Mapping these conformations for SARS-CoV-2 proteins will provide a clearer picture of how they enable the virus to perform diverse functions, such as infecting cells, evading a host’s immune system, and replicating. Such maps may also present new therapeutic opportunities, such as ‘cryptic’ pockets that are absent in experimental snapshots but provide novel targets for drug discovery.

Molecular dynamics simulations have the ability to capture the full ensemble of structures a protein adopts but require significant computational resources. Such simulations capture an all-atom representation of the range of motions a protein undergoes. Modern datasets often consist of a few microseconds of simulation for a single protein, with a few noteworthy examples reaching millisecond timescales.^8,9^ However, many important processes occur on slower timescales. Moreover, simulating every protein that is relevant to SARS-CoV-2 for biologically relevant timescales would require compute resources on an unprecedented scale.

To overcome this challenge, more than a million citizen scientists from around the world have donated their computer resources to simulate SARS-CoV-2 proteins. This massive collaboration was enabled by the Folding@home distributed computing platform, which has crossed the exascale computing barrier and is now the world’s largest supercomputer. Using this resource, we constructed quantitative maps of the structural ensembles of over two dozen proteins and complexes that pertain to SARS-CoV-2. Together, we have run an unprecedented 0.1 s of simulation. Our data uncover the mechanisms of conformational changes that are essential for SARS-CoV-2’s replication cycle and reveal a multitude of new therapeutic opportunities. The data are supported by a variety of experimental observations and are being made publicly available (https://covid.molssi.org/ and https://osf.io/fs2yv/) in accordance with open science principles to accelerate the discovery of new therapeutics.^10,11^

### To the Exascale and beyond!

Folding@home (http://foldingathome.org) is a community of citizen scientists, researchers, and tech organizations dedicated to applying their collective computational and intellectual resources to understand the role of proteins’ dynamics in their function and dysfunction, and to aid in the design of new proteins and therapeutics. The project was founded in the year 2000 with the intent of understanding how proteins fold.^12^ At the time, simulating the folding of even small proteins could easily take thousands of years on a single computer. To overcome this challenge, researchers developed algorithms for dividing these seemingly intractable problems into smaller simulations that could be performed completely independently of one another. They then created the Folding@home project to enable anyone with a computer and an internet connection to volunteer to run these small chunks of simulation, called “work units”.

Over the years, the applications of Folding@home have been generalized to address many aspects of protein dynamics, and the algorithms have developed significantly. The Folding@home Consortium now involves eight laboratories around the world studying various aspects of disease from cancer to antimicrobial resistance to membrane protein dysfunction diseases (https://foldingathome.org/about/the-foldinghome-consortium/). The project has provided insight into diverse topics, ranging from signaling mechanisms.^13-15^ to the connection between phenotype and genotype.^16-18^ Translational applications have included new means to combat antimicrobial resistance, Ebola virus, and SFTS virus.^19-21^

In response to the COVID-19 pandemic, Folding@home quickly pivoted to focus on SARS-CoV-2 and the host factors it interacts with. Many people found the opportunity to take action at a time when they were otherwise feeling helpless alluring. In less than three months, the project grew from ∼30,000 active devices to over a million devices around the globe (Fig. 1A and 1B).

**Figure 1:**
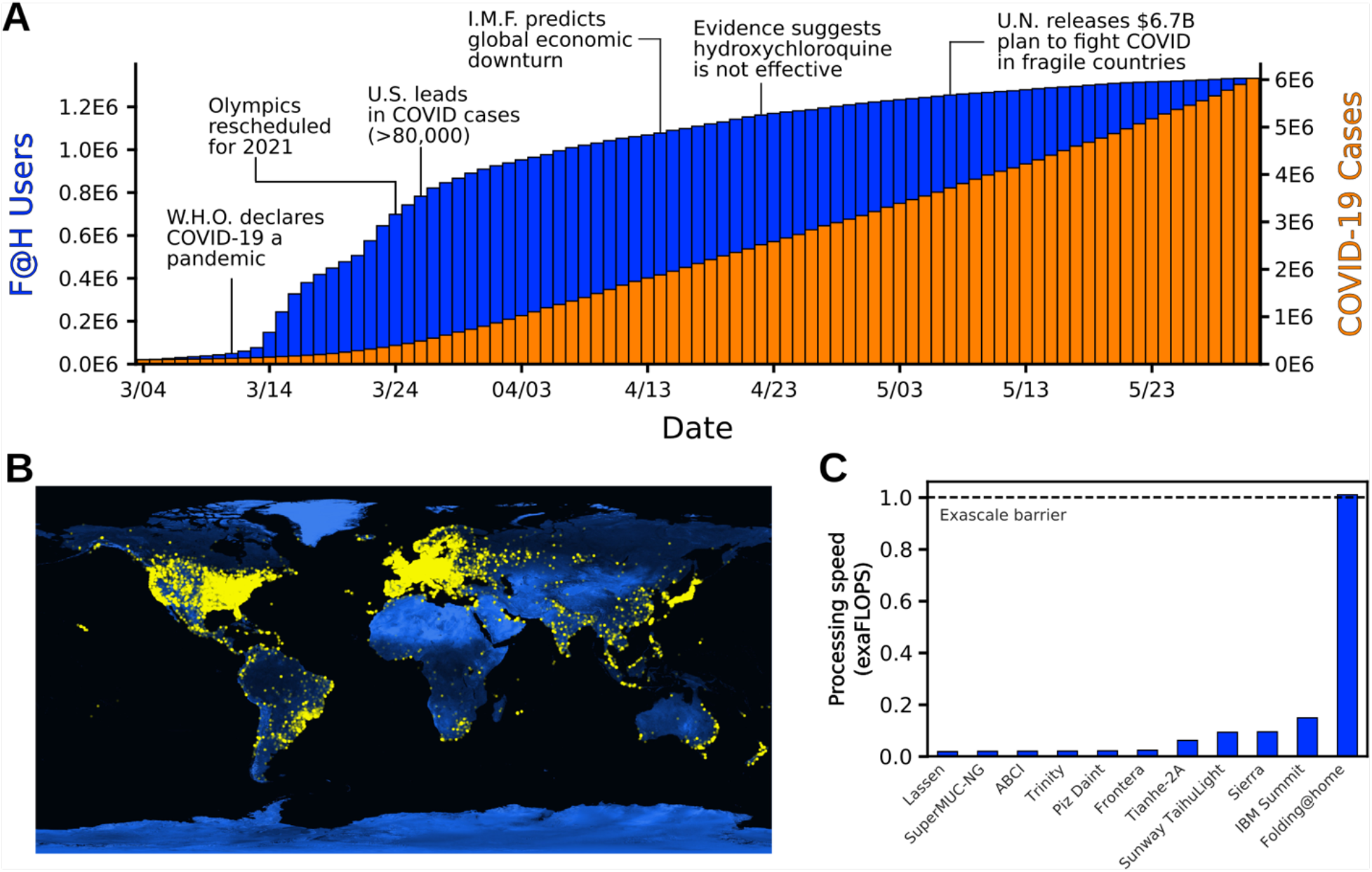
Summary of Folding@home’s computational power. **A)** The growth of Folding@home (F@H) in response to COVID-19. The cumulative number of users is shown in blue and COVID-19 cases are shown in orange. **B)** Global distribution of Folding@home users. Each yellow dot represents a unique IP address contributing to Folding@home. **C)** The processing speed of Folding@home and the next 10 fastest supercomputers, in exaFLOPS.

Estimating the aggregate compute power of Folding@home is non-trivial due to factors like hardware heterogeneity, measures to maintain volunteers’ anonymity, and the fact that volunteers can turn their machines on and off at-will. Furthermore, volunteers’ machines only communicate with the Folding@home servers at the beginning and end of a work unit, with the intervening time taking anywhere from tens of minutes to a few days depending on the volunteer’s hardware and the protein to simulate. Therefore, we chose to estimate the performance by counting the number of GPUs and CPUs that participated in Folding@home during a three-day window and making a conservative assumption about the computational performance of each device (see Methods for details). We note that a larger time window has been used on our website for historical reasons.

Given the above, we conservatively estimate the peak performance of Folding@home hit 1.01 exaFLOPS. This performance was achieved at a point when ∼280,000 GPUs and 4.8 million CPU cores were performing simulations. As explained in the Methods, to be conservative about our claims, we assume that each GPU/CPU has worse performance than a card released before 2015. For reference, the aggregate 1 exaFLOPS performance we report for Folding@home is 5-fold greater than the peak performance of the world’s fastest traditional supercomputer, called Summit (Fig. 1C). It is also more than the top 100 supercomputers combined. Prior to Folding@home, the first exascale supercomputer was not scheduled to come online until the end of 2021.

### Extensive spike opening reveals cryptic epitopes

The Spike complex (S) is a prominent vaccine target that is known to undergo substantial conformational changes as part of its function.^22-24^ Structurally, S is composed of three interlocking proteins, with each chain having a cleavage site separating an S1 and S2 fragment. S resides on the virion surface, where it waits to engage with an angiotensin-converting enzyme 2 (ACE2) receptor on a host cell to trigger infection.^25,26^ The fact that S is exposed on the virion surface makes it an appealing vaccine target. However, it has a number of effective defense strategies. First, S is decorated extensively with glycans that aid in immune evasion by shielding potential antigens.^27,28^ S also uses a conformational masking strategy, wherein it predominantly adopts a closed conformation (often called the down state) that buries the receptor-binding domains (RBDs) to evade immune surveillance mechanisms. To engage with ACE2, S must somehow expose the conserved binding interface of the RBDs. Characterizing the full range of S opening is important for understanding pathogenesis and could provide insights into novel therapeutic options.

To capture S opening, we employed our goal-oriented adaptive sampling algorithm, FAST, in conjunction with Folding@home. The FAST method iterates between running a batch of simulations, building a map called a Markov state model (MSM), ranking the conformational states of this MSM based on how likely starting a new simulation from that state is to yield useful data, and starting a new batch of simulations from the top ranked states.^29,30^ The ranking function is designed to balance between favoring structures with a desired geometric feature (in this case opening of S) and broad exploration of conformational space. By balancing exploration-exploitation tradeoffs, FAST often captures conformational changes with orders of magnitude less simulation time than alternative methods. Broadly distributed structures from our FAST simulations were then used as starting points for extensive Folding@home simulations, totaling over 1 ms of data for SARS-CoV-2 S, enabling us to obtain a statistically sound final model.

Our SARS-CoV-2 S protein simulations capture opening of S and substantial conformational heterogeneity in the open state with full atomistic detail (Fig. 2). Capturing opening of S is an impressive technical feat. Other large-scale simulations have provided valuable insight into aspects of S, but were unable to capture this essential event for the initiation of infection.^28,31,32^ We successfully captured this rare event for both glycosylated and unglycosylated S and found that glycosylation slightly increases the population of the open state, but is qualitatively very similar to the unglycosylated ensemble (Fig. S1). The closed state is more probable than the open state, explaining the experimental observation that full-length S has a lower affinity for ACE2 than an isolated RBD.^40^ Intriguingly, we find that opening occurs only for a single RBD at a time, akin to the up state observed in cryoEM structures.^33^ Moreover, we find that the scale of S opening is often substantially larger than has been observed in experimental snapshots in the absence of binding partners (Fig. S2). The dramatic opening we observe explains the observation that antibodies, and other therapeutics, can bind to regions of the RBD that are deeply buried and seemingly inaccessible in existing experimental snapshots.^34-37^ For example, the cryptic epitope for the antibody CR3022 is buried in up and down cryoEM structures, but is clearly exposed in our conformational ensemble (Fig. 2C). Indeed, our ensemble captures the exposure of many known epitopes, despite their occlusion in apo experimental snapshots (Fig. 2D). Our models also provide a quantitative estimate of the probability that different epitopes are exposed, is consistent with experimental measures of dynamics, and can be used to determine the most suitable regions for the design of neutralizing antibodies.

**Figure 2:**
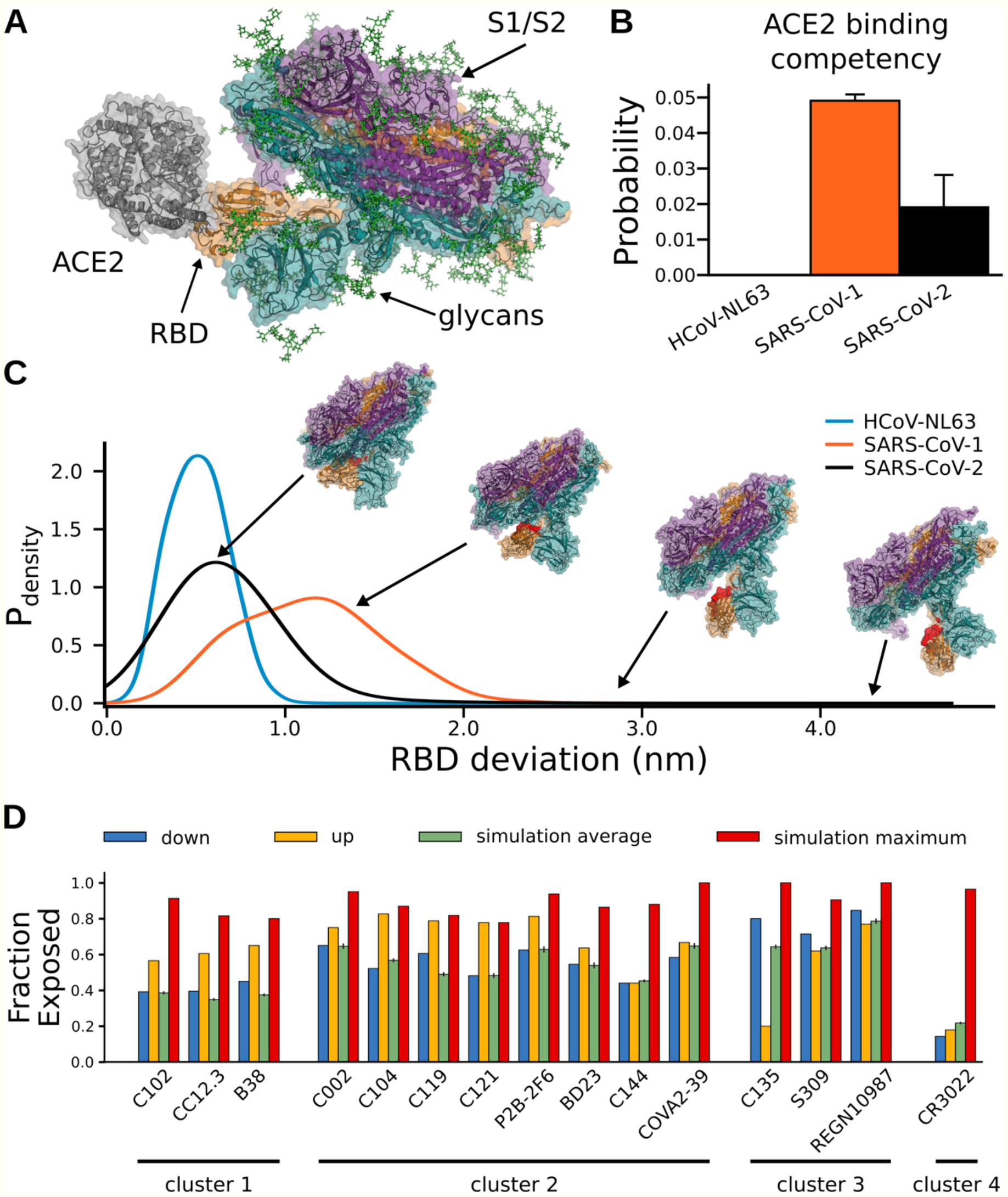
Structural characterization of Spike opening and conformational masking for three Spike homologues. **A)** An example structure of SARS-CoV-2 Spike protein from our simulations that is fully compatible with receptor binding, as shown by superimposing ACE2 (gray). The three chains of Spike are illustrated with a cartoon and transparent surface representation (orange, teal, and purple), and glycans are shown as sticks (green). **B)** Three Spike homologues have very different probabilities of adopting ACE2 binding competent conformations, likely modulating their affinities for both ACE2 and antibodies that engage the ACE2-binding interface. HCoV-NL63, SARS-CoV-1, and SARS-CoV-2 are shown as light-blue, orange, and black, respectively. **C)** The probability distribution of Spike opening for each homologue. Opening is quantified in terms of how far the center of mass of an RBD deviates from its position in the closed (or down) state. The cryptic epitope for the antibody CR3022 (red) is only accessible to antibody binding in extremely open conformations. **D)** Our simulations capture exposure of cryptic epitopes that are buried in the up and down cryoEM structures. The fraction of residues within different epitopes that are exposed to a 0.5 nm radius probe for the down structure (blue), up structure (yellow), the ensemble average from our simulations (green), and the maximum value we observe in our simulations (red). Epitopes are determined as the residues that contact the specified antibody, and are clustered by their binding location on the RBD.^36^

To understand the potential role of conformational masking in determining the lethality and infectivity of different coronaviruses, we also simulated the opening of S proteins from two related viruses: SARS-CoV-1 and HCoV-NL63. These viruses were selected because they also bind the ACE2 receptor but are associated with varying mortality rates. SARS-CoV-1 caused an outbreak in 2003 with a high case fatality rate but has not become a pandemic.^38^ NL63 was discovered the following year and continues to spread around the globe, although it is significantly less lethal than either SARS virus.^39^ We hypothesized that these phenotypic differences may be partially explained by changes to the S conformational ensemble. Specifically, we propose mutations or other perturbations can increase the S-ACE2 affinity by increasing the probability that S adopts an open conformation or by increasing the affinity between an exposed RBD and ACE2. In contrast, the affinity of S for ACE2 (or antibodies that bind cryptic epitopes) can be reduced by stabilizing the closed state.

As expected, the three S complexes have very different propensities to adopt an open state and bind ACE2. Structures from each ensemble were classified as competent to bind ACE2 if superimposing an ACE2-RBD structure on S did not result in any steric clashes between ACE2 and the rest of the S complex. We find that SARS-CoV-1 has the highest population of conformations that can bind to ACE2 without steric clashes, followed by SARS-CoV-2, while opening of NL63 is sufficiently rare that we did not observe ACE2-binding competent conformations in our simulations (Fig. 2B). Interestingly, S proteins that are more likely to adopt structures that are competent to bind ACE2 are also more likely to adopt highly open structures (Fig 2C).

We also observe a number of interesting correlations between conformational masking, lethality, and infectivity of different coronaviruses. First, more deadly coronaviruses have S proteins with less conformational masking. Second, there is an inverse correlation between S opening and the affinity of an isolated RBD for ACE2 (RBD-ACE2 affinities of ∼35 nM, ∼44 nM, and ∼185 nM for HCoV-NL63, SARS-CoV-2, and SARS-CoV-1, respectively).^41,42^

These observations suggest a tradeoff wherein greater conformational masking enables immune evasion but requires a higher affinity between an exposed RBD and ACE2 to successfully infect a host cell. We propose that the NL63 S complex is probably best at evading immune detection but is not as infectious as the SARS viruses because strong conformational masking reduces the overall affinity for ACE2. In contrast, the SARS-CoV-1 S complex adopts open conformations more readily but is also more readily detected by immune surveillance mechanisms. Finally, SARS-CoV-2 balances conformational masking and the RBD-ACE2 affinity in a manner that allows it to evade an immune response while maintaining its ability to infect a host cell. Based on this model, we predict that mutations that increase the probability that the SARS-CoV-2 S complex adopts open conformations may be more lethal but spread less readily.

Our atomically detailed model of S can facilitate structure-based vaccine antigen design through identification of regions minimally protected by conformational masking or the glycan shield.^43^ To identify these potential epitopes, we calculated the probability that each residue in S could be exposed to therapeutics (e.g. not shielded by a glycan or buried by conformational masking), as shown in Fig. 3A. Visualizing these values on the protein reveals a few patches of protein surface that are exposed through the glycan shielding (Fig. 3B). However, another important factor when targeting an antigen is picking a region with a conserved sequence to yield broader and longer lasting efficacy. Not surprisingly, many of the exposed regions do not have a strongly conserved sequence. Promisingly, though, we do find a conserved area with a larger degree of solvent exposure (Fig. 3C). This region was recently found to be an effective site for neutralizing antibodies.^44^ Another possibility for antigen design is to exploit the opening motion. A number of residues surrounding the receptor binding motif (RBM) of the RBD show an increase in exposure by ∼30% in ACE2 binding competent structures (Fig. 3C). Consistent with immunoassays and cryoEM structures, these regions are hotspots for neutralizing antibody binding.^34,45,46^

**Figure 3:**
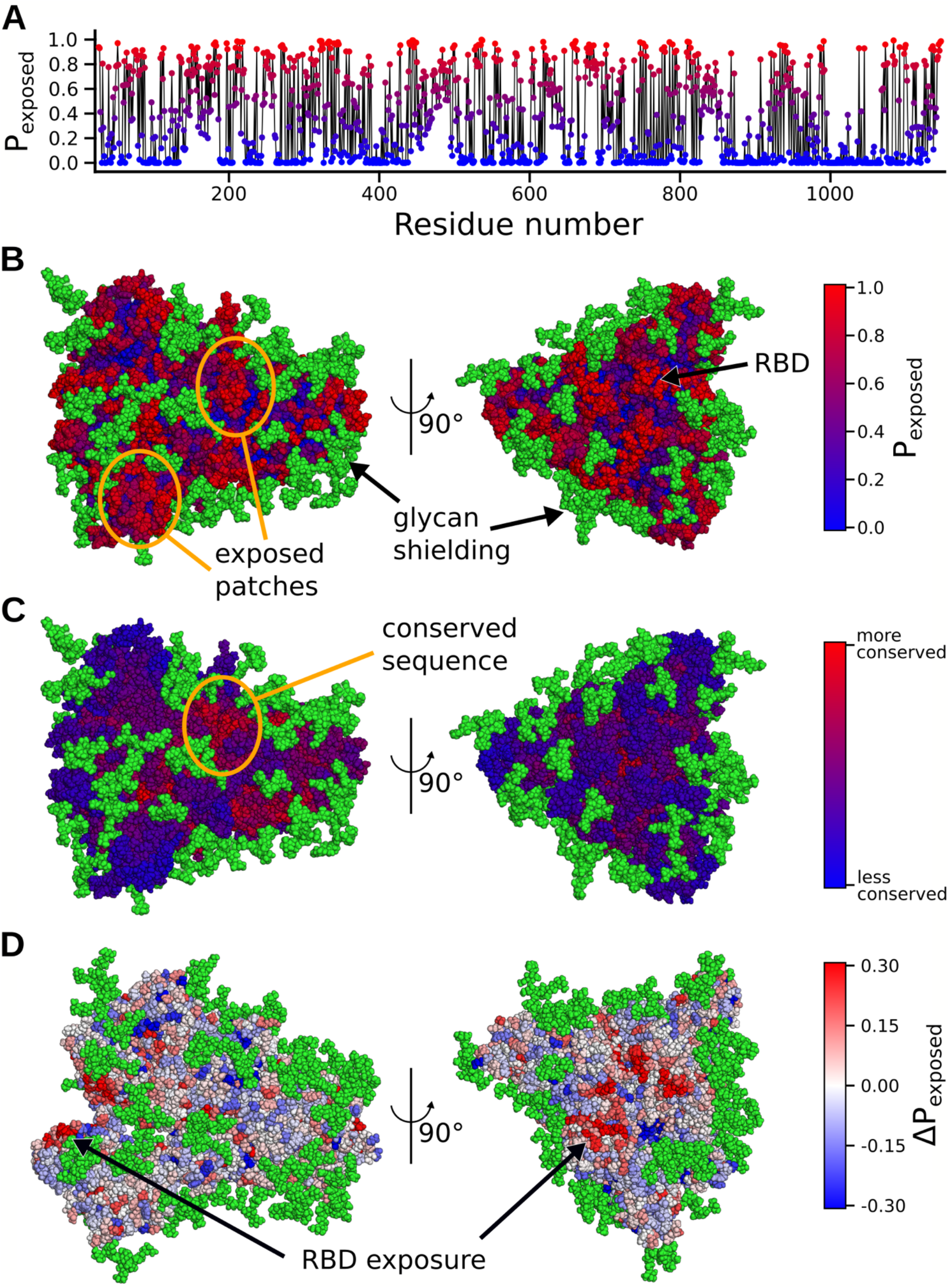
Effects of glycan shielding and conformational masking on the accessibility of different parts of the Spike to potential therapeutics. **A)** The probability that a residue is exposed to potential therapeutics, as determined from our structural ensemble. Red indicates a high probability of being exposed and blue indicates a low probability of being exposed. **B)** Exposure probabilities colored on the surface of the Spike protein. Exposed patches are circled in orange. Red residues have a higher probability of being exposed, whereas blue residues have a lower probability of being exposed. Green atoms denote glycans. **C)** Sequence conservation score colored onto the Spike protein. A conserved patch on the protein is circled in orange. Red residues have higher conservation, whereas blue residues have lower conservation. **D)** The difference in the probability that each residue is exposed between the ACE2-binding competent conformations and the entire ensemble. Red residues have a higher probability of being exposed upon opening, whereas blue residues have a lower probability of being exposed.

### Cryptic pockets and functional dynamics are present throughout the proteome

Every protein in SARS-CoV-2 remains a potential drug target. So, to understand their role in disease and help progress the design of antivirals, we unleashed the full power of Folding@home to simulate dozens of systems related to pathogenesis. While we are interested in all aspects of a proteins’ functional dynamics, expanding on the number of antiviral targets is of immediate value. Towards this end, we seeded Folding@home simulations from our FAST-pockets adaptive sampling to aid in the discovery of cryptic pockets. We briefly discuss two illustrative examples, out of 36 datasets.

Nonstructural protein number 5 (NSP5, also named the main protease, 3CL^pro^, or as we will refer to it, M^pro^) is an essential protein in the lifecycle of coronaviruses, cleaving polyprotein 1a into functional proteins, and is a major target for the design of antivirals.^47^ It is highly conserved between coronaviruses and shares 96% sequence identity with SARS-CoV-1 M^pro^; it cleaves polyprotein 1a at no fewer than 11 distinct sites, placing significant evolutionary constraint on its active site. M^pro^ is only active as a dimer, however it exists in a monomer-dimer equilibrium with estimates of its dissociation constant in the low μM range.^48^ Small molecules targeting this protein to inhibit enzymatic activity, either by altering its active site or favoring the inactive monomer state, would be promising broad-spectrum antiviral candidates.^49^

Our simulations reveal two novel cryptic pockets on M^pro^ that expand our current therapeutic options. These are shown in Fig. 4A, which projects states from our MSM onto the solvent exposure of residues that make up the pockets. The first cryptic pocket is an expansion of NSP5’s catalytic site. We find that the loop bridging domains II and III is highly dynamic and can fully undock from the rest of the protein. This motion may impact catalysis—i.e. by sterically regulating substrate binding—and is similar to motions we have observed previously for the enzyme β-lactamase.^50^ Owing to its location, a small molecule bound in this pocket is likely to prevent catalysis by obstructing polypeptide association with catalytic residues. The second pocket is a large opening between domains I/II and domain III. Located at the dimerization interface, this pocket offers the possibility to find small molecules or peptides that favor the inactive monomer state.

**Figure 4:**
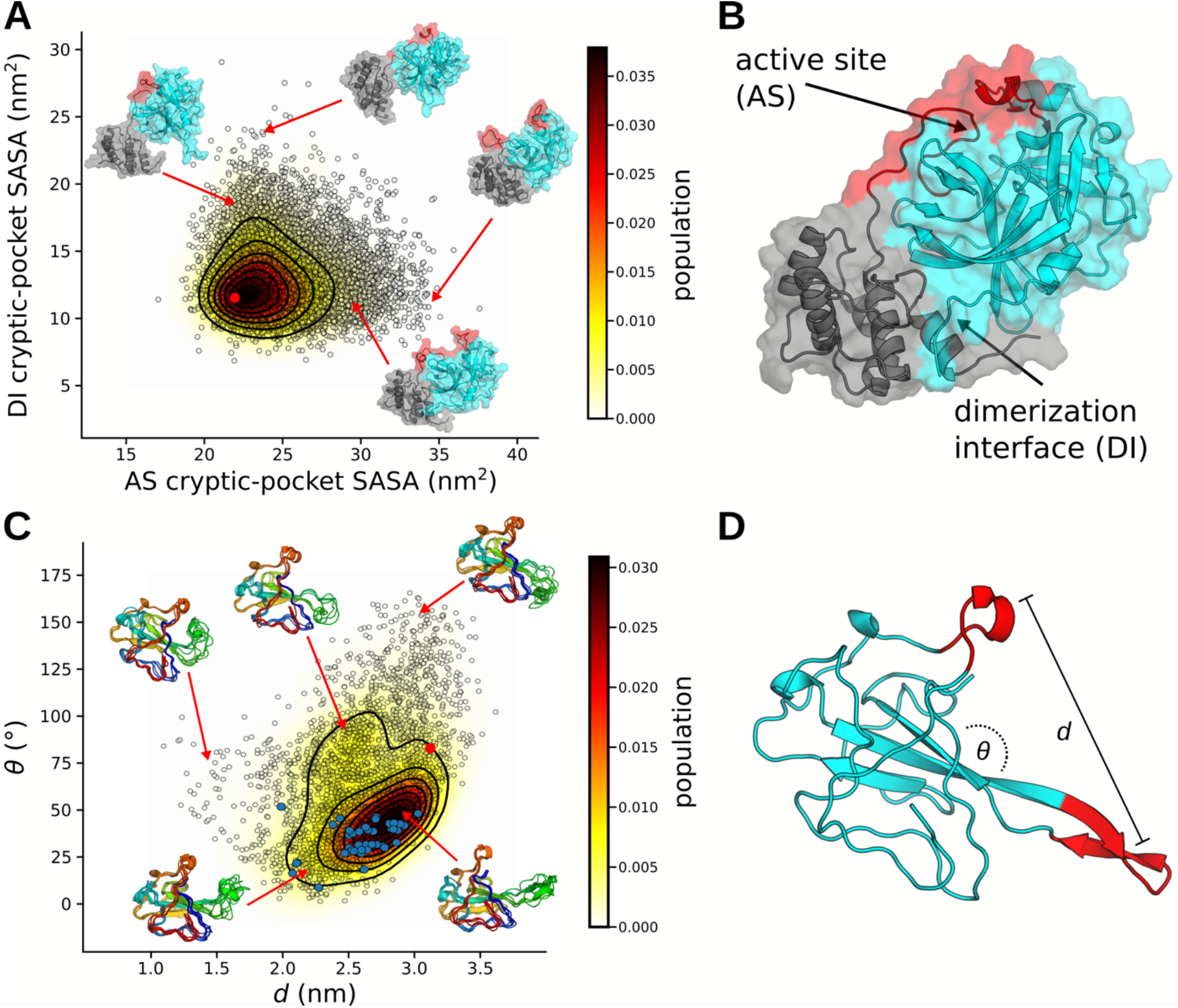
Examples of cryptic pockets and functionally-relevant dynamics. **A-B)** Conformational ensemble of M^pro^ (monomeric) highlighting cryptic pockets near the active site (AS) and domain interface (DI). Conformational states (black circles) are projected onto the solvent accessible surface areas (SASAs) of residues surrounding either the active-site or dimerization interface. The starting structure for simulations (6Y2E) is shown as a red dot. Representative structures are depicted with cartoon and transparent surface. Domains I and II are colored cyan and domain III is colored gray. The loop of domain III, which covers the active-site residues and is seen to be highly dynamic, is colored red. **C-D)** The conformational ensemble from our simulations of nucleoprotein is similar to the distribution of structures seen experimentally. Conformational states are projected onto the distance and angle between the positive finger and a nearby loop. Angles were calculated between vectors that point along each red segment in panel D and distances were calculated between their centers of mass. Cluster centers are represented as black circles, the starting structure for simulations (6VYO) is shown as a red dot, and NMR structures are shown with solid blue dots. Representative structures are shown as cartoons.

**Figure 5:**
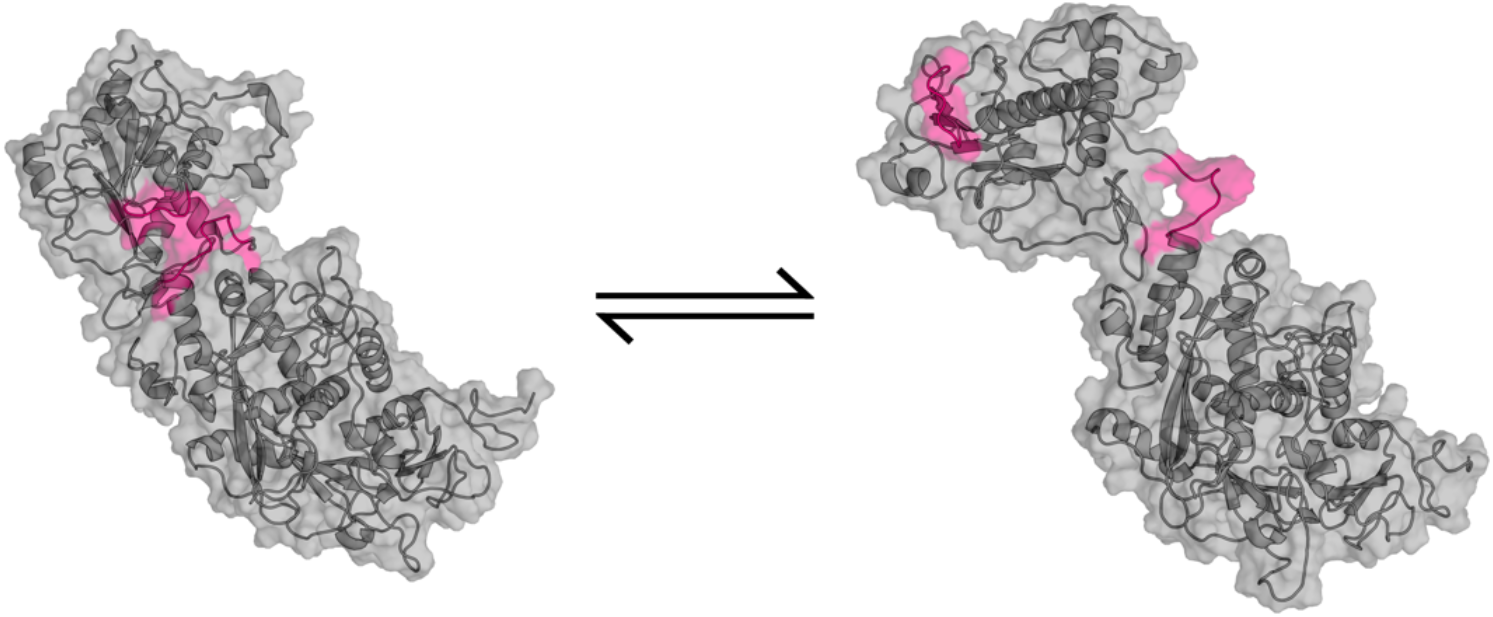
NSP10/NSP14 (complex) transition from closed to open state. Backbone is represented as a cartoon and sidechains are represented with a transparent surface. The residues that undergo a large conformational change to expose a cryptic pocket are highlighted in pink.

**Figure 6:**
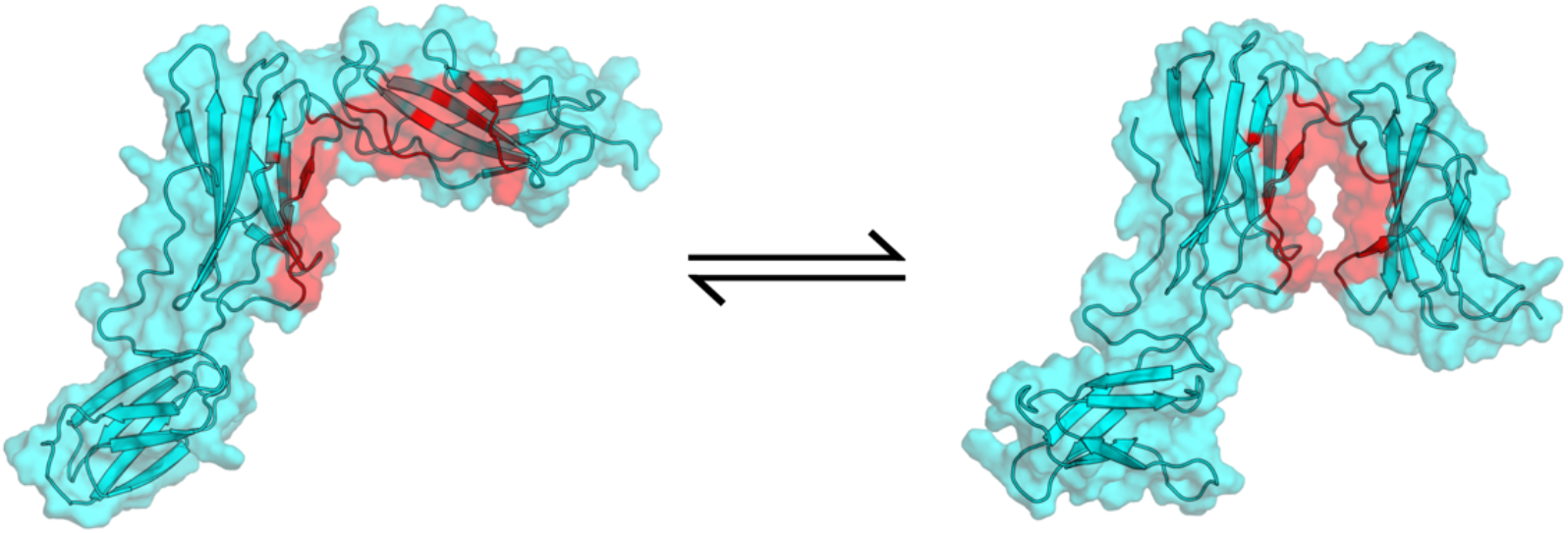
Human IL6-R transition from expanded to closed state. Backbone is represented as a cartoon and sidechains are represented with a transparent surface. The residues that undergo a large conformational change to reveal a potential druggable site are highlighted in red.

In addition to cryptic pockets, our data captures many potentially functionally relevant motions within the SARS-CoV-2 proteome. We illustrate this with the SARS-CoV-2 nucleoprotein. The nucleoprotein is a multifunctional protein responsible for major lifecycle events such as viral packaging, transcription, and physically linking RNA to the envelope.^51,52^ As such, we expect the protein to accomplish these goals through a highly dynamic and rich conformational ensemble, akin to context-dependent regulatory modules observed in Ebola virus nucleoprotein.^53,54^ Investigating the RNA-binding domain, we observe both cryptic pockets and an incredibly dynamic beta-hairpin, which hosts the RNA binding site, referred to as a “positive finger” (Fig. 4C-D). Our observed conformational heterogeneity of the positive finger is consistent with a structural ensemble determined using solution-state nuclear magnetic resonance (NMR) spectroscopy.^55^ Our simulations also capture numerous states of the putative RNA binding pose, where the positive finger curls up to form a cradle for RNA. These states can provide a structural basis for the design of small molecules that would compete with RNA binding, preventing viral assembly.

The data we present in this paper represents the single largest collection of all-atom simulations. Table 1 is a comprehensive list of the systems we have simulated. Systems span various oligomerization states, include important complexes, and include representation from multiple coronaviruses. We also include human proteins that are targets for supportive therapies and preventative treatments. To accelerate the discovery of new therapeutics and promote open science, our MSMs and structures of cryptic pockets are available online (https://covid.molssi.org/ and https://osf.io/fs2yv/). For ease of use, within each final model we provide the residues that comprise cryptic pockets along with an ordered list of states from largest to smallest opening.

**Table 1:** Summary of protein systems we have simulated on Folding@home, organized by viral strain. *Missing residues were modeled using Swiss model.^56^ **Structural model was generated from a homologous sequence using Swiss model.^56^ ***Missing residues were modeled using CHARMM-GUI.^57,58^

| System name | Oligomerization | Initial structure | Residues | Atoms in system | Aggregate simulation time (μs) | Cryptic pockets discovered |
| --- | --- | --- | --- | --- | --- | --- |
| **SARS-CoV-2** |  |  |  |  |  |  |
| NSP3 (Macrodomain “X”) | Monomer | 6W02 | 167 | 23907 | 10,906 | - |
| NSP3 (Papain-like protease 2, PL2<sup>pro</sup>) | Monomer | 3E9S** | 306 | 97285 | 731 | 2 |
| NSP5 (main protease, M<sup>pro</sup>, 3CL<sup>pro</sup>) | Monomer | 6Y2E | 306 | 64791 | 6,405 | 2 |
| NSP5 (main protease, M<sup>pro</sup>, 3CL<sup>pro</sup>) | Dimer | 6Y2E | 612 | 77331 | 2,902 | 2 |
| NSP7 | Monomer | 5F22** | 79 | 20094 | 3,722 | 3 |
| NSP8 | Monomer | 2AHM** | 191 | 156282 | 1,776 | 3 |
| NSP9 | Dimer | 6W4B* | 226 | 49885 | 8,939 | 2 |
| NSP10 | Monomer | 6W4H* | 131 | 29560 | 6,141 | 2 |
| NSP12 (polymerase) | Monomer | 6NUR** | 891 | 186622 | 3,330 | 3 |
| NSP13 (helicase) | Monomer | 6JYT** | 596 | 129368 | 3,407 | 3 |
| NSP14 | Monomer | 5C8S** | 527 | 216380 | 2,384 | 2 |
| NSP15 | Monomer | 6VWW | 347 | 67345 | 3,674 | 4 |
| NSP15 | Hexamer | 6VWW | 2082 | 230339 | 4,270 | - |
| NSP16 | Monomer | 6W4H* | 298 | 45672 | 2,382 | 5 |
| Nucleoprotein (RBD) | Monomer | 6VYO | 173 | 29125 | 9,493 | 3 |
| Nucleoprotein Dimerization Domain | Monomer | 6YUN* | 118 | 34905 | 6,782 | - |
| Nucleoprotein Dimerization Domain | Dimer | 6YUN* | 236 | 72733 | 1,458 | 2 |
| Spike | Trimer | 6VXX*** | 3363 | 442881 | 1,109 | - |
| NSP7 / NSP8 / NSP12 | Trimer complex | 6NUR** | 1184 | 215694 | 1,686 | - |
| NSP10 / NSP14 | Dimer complex | 5C8S** | 688 | 226672 | 689 | 3 |
| NSP10 / NSP16 | Dimer complex | 6W4H* | 429 | 63752 | 3,463 | 2 |
| **SARS-CoV-1** |  |  |  |  |  |  |
| NSP3 (Macrodomain “X”) | Monomer | 2FAV | 172 | 33117 | 659 | - |
| NSP9 | Dimer | 1QZ8* | 226 | 49599 | 7,763 | - |
| NSP15 | Monomer | 2H85 | 345 | 67345 | 4,734 | - |
| NSP15 | Hexamer | 2H85 | 2070 | 230339 | 1,130 | - |
| Nucleoprotein RBD | Monomer | 2OFZ | 174 | 29125 | 4,088 | - |
| Nucleoprotein Dimerization Domain | Monomer | 2GIB | 370 | 34905 | 1,626 | - |
| Nucleoprotein Dimerization Domain | Dimer | 2GIB | 740 | 72733 | 4,221 | - |
| Spike | Trimer | 5X58*** | 3261 | 375851 | 741 | - |
| NSP10 / NSP16 | Dimer complex | 6W4H** | 425 | 69589 | 518 | - |
| **Human** |  |  |  |  |  |  |
| IL6 | Monomer | 1ALU | 166 | 26855 | 1,593 | 2 |
| IL6-R | Monomer | 1N26 | 299 | 149764 | 196 | 5 |
| ACE2 | Monomer | 6LZG | 596 | 75787 | 664 | 2 |
| **MERS** |  |  |  |  |  |  |
| NSP13 | Monomer | 5WWP | 596 | 121134 | 719 | - |
| NSP10 / NSP16 | Dimer Complex | 6W4H** | 424 | 69127 | 518 | - |
| **HCoV-NL63** |  |  |  |  |  |  |
| Spike | Trimer | 5SZS*** | 3606 | 453348 | 651 | - |

## Discussion

The Folding@home community has created one of the largest computational resources in the world to tackle a global threat. Over a million citizen scientists have pooled their computer resources to help understand and combat COVID-19, generating more than 0.1 seconds of simulation data. The unprecedented scale of these simulations has helped to characterize crucial stages of infection. We find that Spike proteins have a strong trade-off between making ACE2 binding interfaces accessible to infiltrate cells and conformationally masking epitopes to subvert immune responses. SARS-CoV-2 represents a more optimal tradeoff than related coronaviruses, which may explain its success in spreading globally. Our simulations also provide an atomically detailed roadmap for designing vaccines and antivirals. For example, we have made a comprehensive atlas and repository of cryptic pockets hosted online to accelerate the development of novel therapeutics. Many groups are already using our data, such as the COVID Moonshot,^59^ an international collaboration between multiple computational and experimental groups working to develop a patent-free inhibitor of the main protease.

Beyond SAR S-CoV-2, we expect this work to aid in a better understanding of the roles of proteins in the *coronaviridae* family. Coronaviruses have been around for millennia, yet many of their proteins are still poorly understood. Because climate change has made zoonotic transmission events more commonplace, it is imperative that we continue to perform basic research on these viruses to better protect us from future pandemics. For each protein system in Table 1, an extraordinary amount of sampling has led to the generation of a quantitative map of its conformational landscape. There is still much to learn about coronavirus function and these conformational ensembles contain a wealth of information to pull from.

While we have aggressively targeted research on SARS-CoV-2, Folding@home is a general platform for running molecular dynamics simulations at scale. Before the COVID-19 pandemic, Folding@home was already generating datasets that were orders of magnitude greater than from conventional means. With our explosive growth, our compute power has increased around 100-fold. Our work here highlights the incredible utility this compute power has to rapidly understand health and disease, providing a rich source of structural data for accelerating the design of therapeutics. With the continued support of the citizen scientists that have made this work possible, we have the opportunity to make a profound impact on other global health crises such as cancer, neurodegenerative diseases, and antibiotic resistance.

## Methods

### System preparation

All simulations were prepared using Gromacs 2020.^60^ Initial structures were placed in a dodecahedral box that extends 1.0 nm beyond the protein in any dimension. Systems were then solvated and energy minimized with a steepest descents algorithm until the maximum force fell below 100 kJ/mol/nm using a step size of 0.01 nm and a cutoff distance of 1.2 nm for the neighbor list, Coulomb interactions, and van der Waals interactions. The AMBER03 force field was used for all systems except Spike protein with glycans, which used CHARMM36.^61,62^ All simulations were simulated with explicit TIP3P solvent.^63^

Systems were then equilibrated for 1.0 ns, where all bonds were constrained with the LINCS algorithm and virtual sites were used to allow a 4 fs time step.^64^ Cutoffs of 1.1 nm were used for the neighbor list with 0.9 for Coulomb and van der Waals interactions. The particle mesh ewald method was employed for treatment of long-range interactions with a fourier spacing of 0.12 nm. The Verlet cutoff scheme was used for the neighbor list. The stochastic velocity rescaling (*v*-rescale) thermostat was used to hold the temperature at 300 K.^65^

### Adaptive sampling simulations

The FAST algorithm was employed for each protein in Table 1 to enhance conformational sampling and quickly explore dominant motions. The procedure for FAST simulations is as follows: 1) run initial simulations, 2) build MSM, 3) rank states based on FAST ranking, 4) restart simulations from the top ranked states, 5) repeat steps 2-4 until ranking is optimized. For each system, MSMs were generated after each round of sampling using a *k*-centers clustering algorithm based on the RMSD between select atoms. Clustering continued until the maximum distance of a frame to a cluster center fell within a predefined cutoff. In addition to the FAST ranking, a similarity penalty was added to promote conformational diversity in starting structures, as has been described previously.^66^

FAST-distance simulations of all Spike proteins were run at 310 K on the Microsoft Azure cloud computing platform. The FAST-distance ranking favored states with greater RBD openings using a set of distances between atoms. Each round of sampling was performed with 22 independent simulations that were 40 ns in length (0.88 μs aggregate sampling per round), where the number of rounds totaled 13 (11.44 μs), 22 (19.36 μs), and 17 (14.96 μs), for SARS-CoV-1, SARS-CoV-2, and HCoV-NL63, respectively.

For all other proteins, FAST-pocket simulations were run at 300 K for 6 rounds, with 10 simulations per round, where each simulation was 40 ns in length (2.4 μs aggregate simulation). The FAST-pocket ranking function favored restarting simulations from states with large pocket openings. Pocket volumes were calculated using the LIGSITE algorithm.^67^

### Folding@home simulations

For each adaptive sampling run, a conformationally diverse set of structures was selected to be run on Folding@home. Structures came from the final *k*-centers clustering of adaptive sampling, as is described above. Simulations were deployed using a simulation core based on either GROMACS 5.0.4 or OpenMM 7.4.1.^60,68^

To estimate the performance of Folding@home, we make the conservative assumption that each CPU core performs at 0.0127 TFLOPS and each GPU at 1.672 native TFLOPS (or 3.53 X86-equivalent TFLOPS), as explained in our long-standing performance estimate (https://stats.foldingathome.org/os). For reference, a GTX 980 (which was released in 2014) can achieve 5 native TFLOPS (or 10.56 X86-equivalent TFLOPS). An Intel Core i7 4770K (released in 2013) can achieve 0.046 TFLOPS/core. We report x86-equivalent FLOPS.

### Markov state models

A Markov state model is a network representation of a free energy landscape and is a key tool for making sense of molecular dynamics simulations.^69^ All MSMs were built using our python package, enspara.^70^ Each system was clustered with the combined FAST and Folding@home datasets. In the case of Spike proteins, states were defined geometrically based on the RMSD between backbone C_α_ coordinates. States were generated as the top 3000 centers from a *k*-centers clustering algorithm. All other proteins were clustered based on the Euclidean distance between the solvent accessible surface area of residues, as is described previously.^50^ Systems generated either 2500, 5000, 7500, or 10000 cluster centers from a *k*-centers clustering algorithm. Select systems were refined with 1-10 *k*-medoid sweeps. Transition probability matrices were produced by counting transitions between states, adding a prior count of 1/*n*_*states*_, and row-normalizing, as is described previously.^71^ Equilibrium populations were calculated as the eigenvector of the transition probability matrix with an eigenvalue of one.

### Spike/ACE2 binding competency

To determine Spike protein binding competency to ACE2 the following structures of the RBD bound to ACE2 were used: 3D0G, 6M0J, and 3KBH, for SARS-CoV-1, SARS-CoV-2, and HCoV-NL63, respectively. The RBD of the bound complex was superimposed onto each RBD for structures in our MSM. Steric clashes were then determined between backbone atoms on the ACE2 molecule and the rest of the spike protein. If any of the structures had a superposition that resulted in no clashes, it was deemed binding competent.

### Cryptic pockets and solvent accessible surface area

For ease of detecting cryptic pockets and other functional motions, we employed our exposon analysis method.^50^ This method correlates the solvent exposure between residues to find concerted motions that tend to represent cryptic pocket openings. Solvent accessible surface area calculations were computed using the Shrake-Rupley algorithm as implemented in the python package MDTraj.^72^ For all proteins and complexes, a solvent probe radius of 0.28 nm was used, which has been shown to produce a reasonable clustering and exposon map.^50^

Spike protein solvent accessible surface areas for SARS-CoV-2 were computed with glycan chains modeled onto each cluster center. Multiple glycan rotamers were sampled for each state and accessible surface areas for each residue were weighted based on MSM equilibrium populations.

### Sequence conservation

Sequence conservation of spike proteins was calculated using the Uniprot database.^73^ Sequences between 30% - 90% were pulled and aligned with the Muscle algorithm.^74^ The entropy at each position was calculated to quantify variability of amino acids. Conservation was defined as one minus the entropy.

## Acknowledgements

We are extremely grateful to all the citizen scientists who contributed their compute power to make this work possible, and members of the Folding@home community who volunteered to help with everything from technical support to translating content into multiple languages. Thanks to Microsoft AI for Health for helping us use Azure to run adaptive sampling simulations, and to UKRI for providing compute resources to parallelize data analysis. Thanks to Pure Storage for providing a FlashBlade system to store our large datasets, to Seagate and Micron for additional storage, and to MolSSI for helping organize public datasets. Thanks to Avast, AWS, Cisco, Linus Tech Tips, Microsoft Azure, Oracle, and VMware for helping us to scale-up Folding@home’s server-side infrastructure to keep up with the tremendous growth we experienced in such a short time. Thanks to AMD, ARM and Neocortix, and Intel for helping to improve the performance of Folding@home on their hardware. Thanks to all of these companies for helping to spread the word about Folding@home, and also to A16Z, Best Buy, CCP, CoreWeave, Daimler Truck AG, Dell, GitHub, HP, La Liga, Media Monks, Microcenter, NVIDIA, and Telefonica. Thanks to CERN and the particle physics community for helping with data management and to DataDog for server monitoring services. Thanks to Christopher O. Barnes for providing the epitope contacts used for Fig. 2D. JRP acknowledges support from F30HL146052. GRB and his lab were supported by funding from Avast, the Center for the Science and Engineering of Living Systems (CSELS), an NSF RAPID award, NSF CAREER Award MCB-1552471, NIH R01 GM124007, a Burroughs Wellcome Fund Career Award at the Scientific Interface, and a Packard Fellowship for Science and Engineering. JDC acknowledges support from NIH grant P30 CA008748 and NIH grant R01 GM121505. VAV and MFDH acknowledge support from NIH grant R01 GM123296, NIH grant S10-OD020095, and NSF MRI grant CNS-1625061.

## Supporting Information

**Figure S1:**
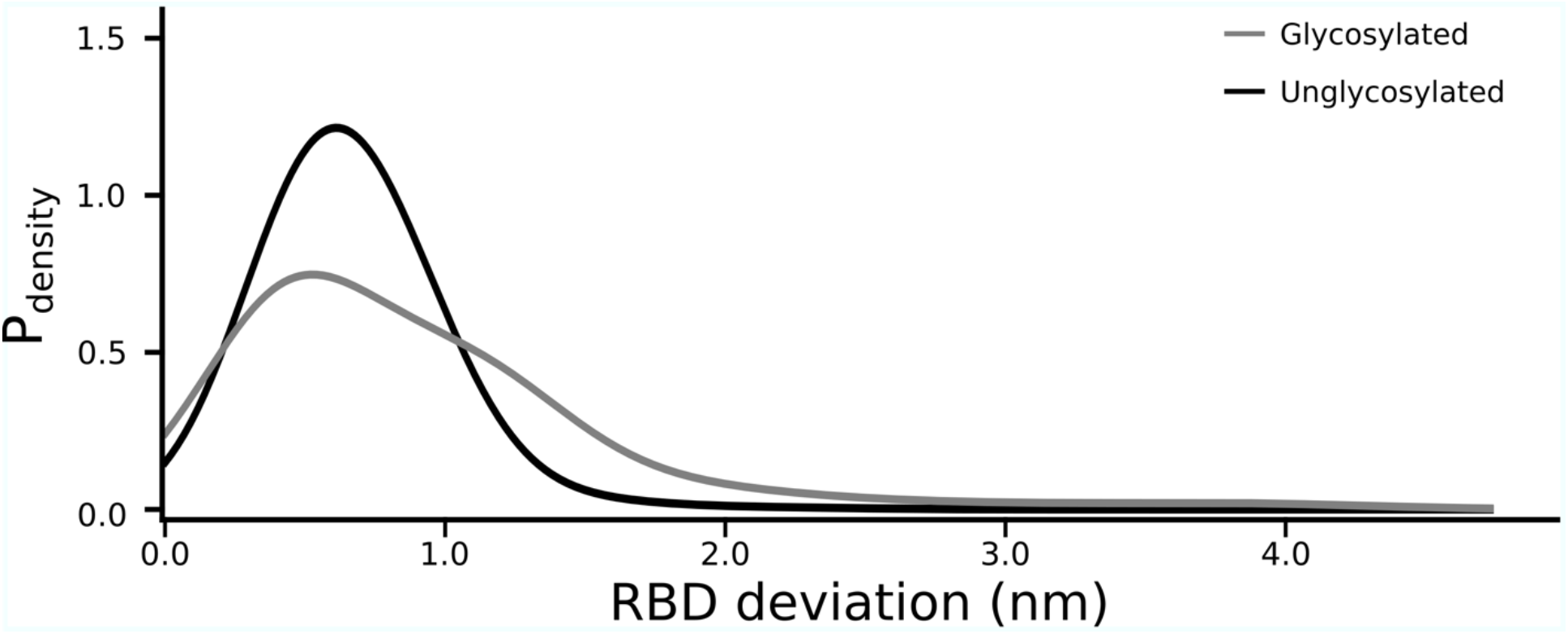
Distribution of SARS-CoV-2 Spike RBD opening. The probability that the center of mass of an RBD deviates from its position in the closed (or down) state for SARS-CoV-2 spike with glycans (gray) and without glycans (black).

**Figure S2:**
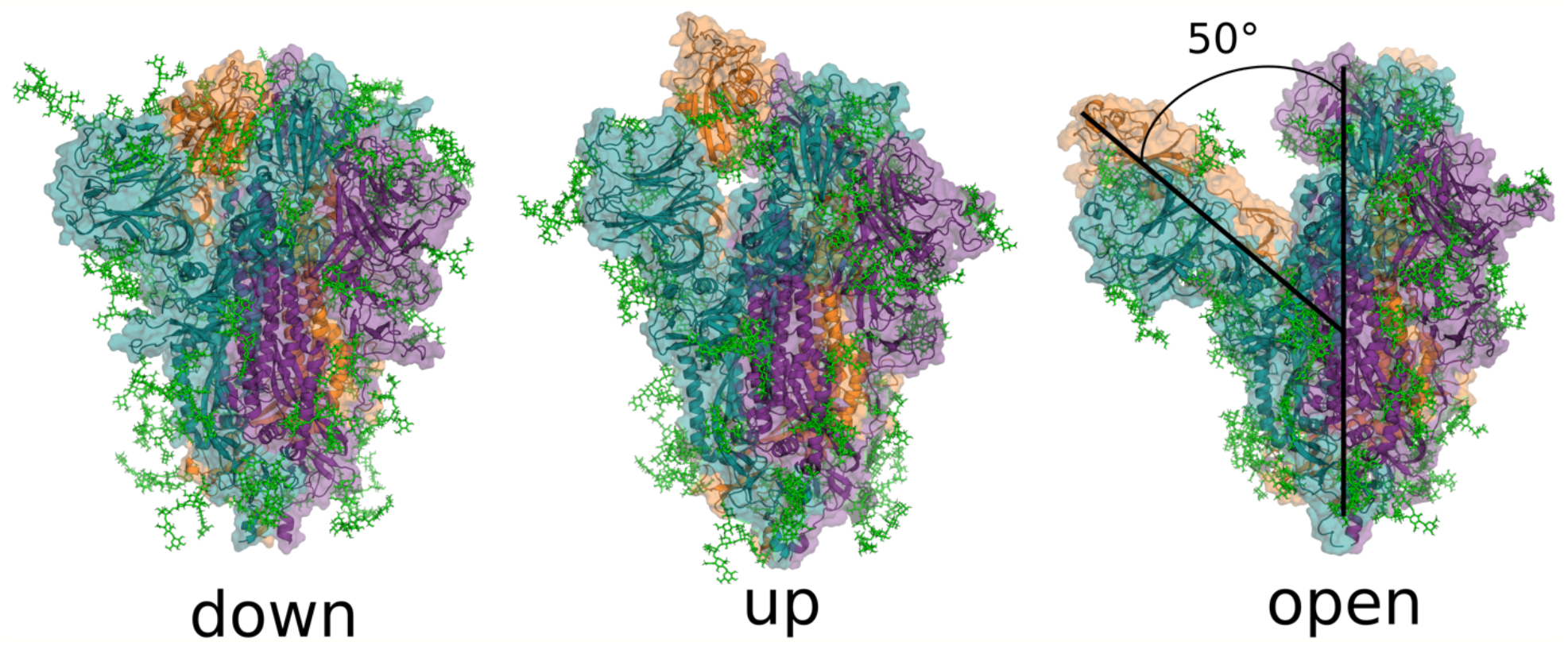
Simulations of the SARS-CoV-2 Spike complex reveal the existence of an “open” state. For reference, three Spike complex snapshots are shown: the “down” state (6VXX), the “up” state (6VSB), and an “open” state from our simulations. Structures are depicted with a cartoon backbone, transparent surface for sidechains, and sticks for glycans. Each chain in the complex has a unique color, orange, purple, or teal, and glycans are colored green.

### Cryptic pocket highlights for select systems in the SARS-CoV-2 proteome

**Figure S3:**
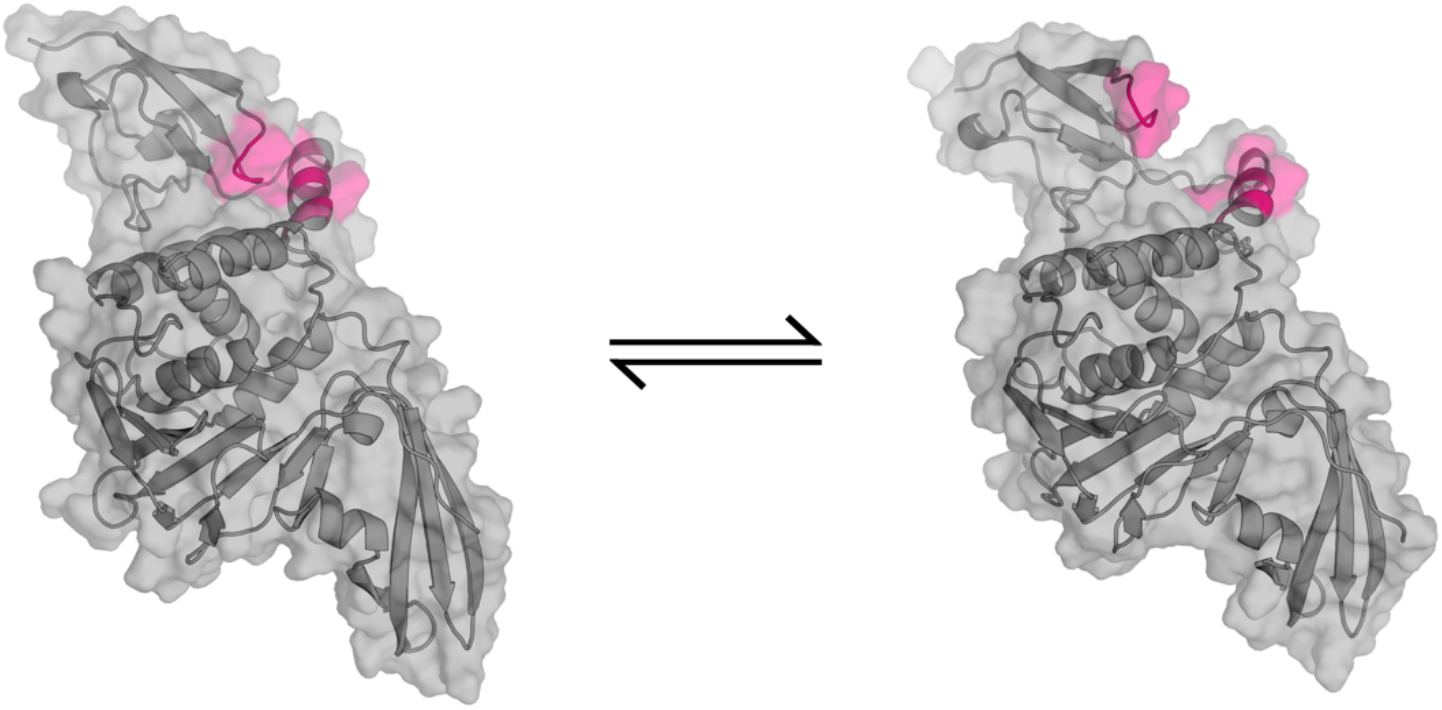
NSP3-PL2Pro domain transition from closed to open state. Backbone is represented as a cartoon and sidechains are represented with a transparent surface (gray). The residues that undergo a large conformational change to expose a cryptic pocket are highlighted in pink.

**Figure S4:**
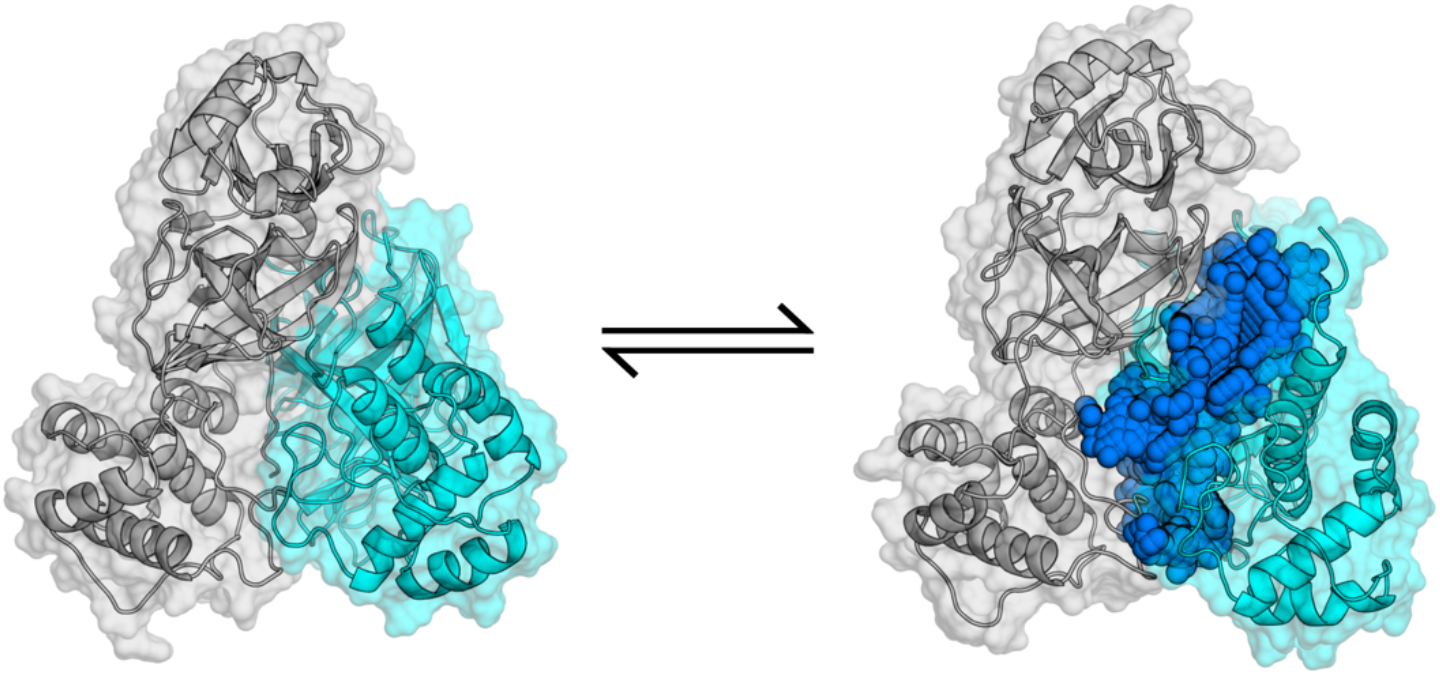
NSP5 (dimer) transition from closed to open state. Backbone is represented as a cartoon, sidechains are represented with a transparent surface, and pocket volumes are represented as blue spheres. Each molecule in the dimer is identified with a unique color, gray or cyan.

**Figure S5:**
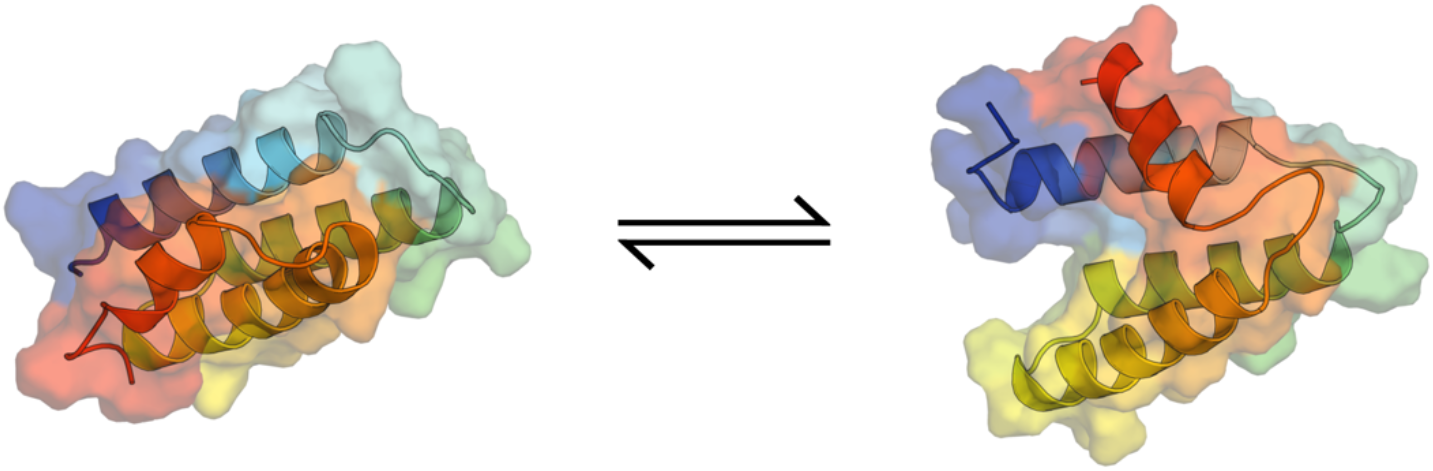
NSP7 transition from closed to open state. Backbone is represented as a cartoon and sidechains are represented with a transparent surface. The protein is colored by residue number following a rainbow.

**Figure S6:**
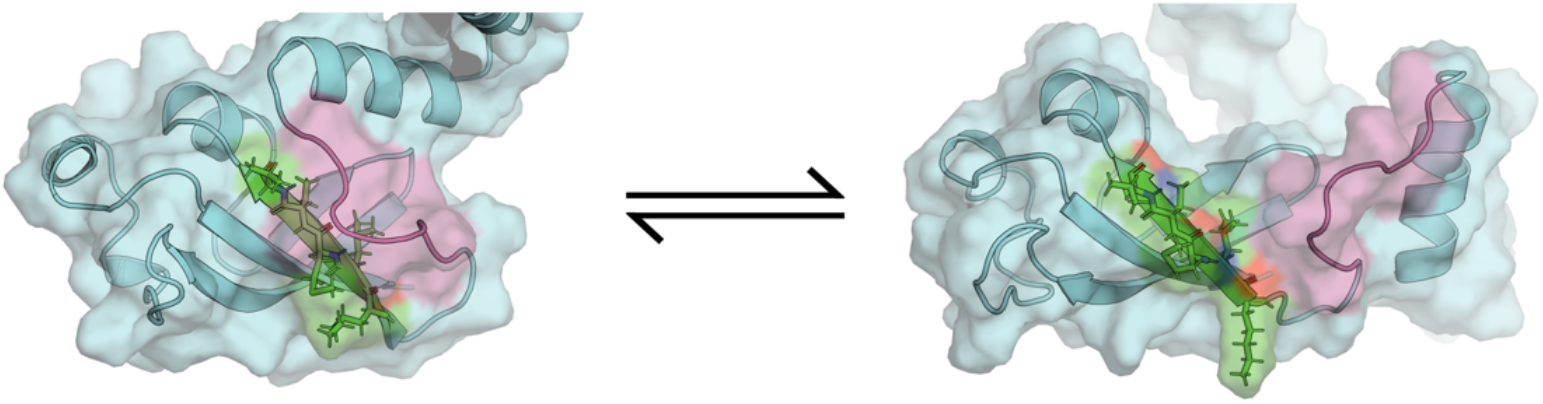
NSP8 transition from closed to open state. Backbone is represented as a cartoon and sidechains are represented with a transparent surface. For reference, two regions that undergo a large conformational transition are highlighted as green and pink.

**Figure S7:**
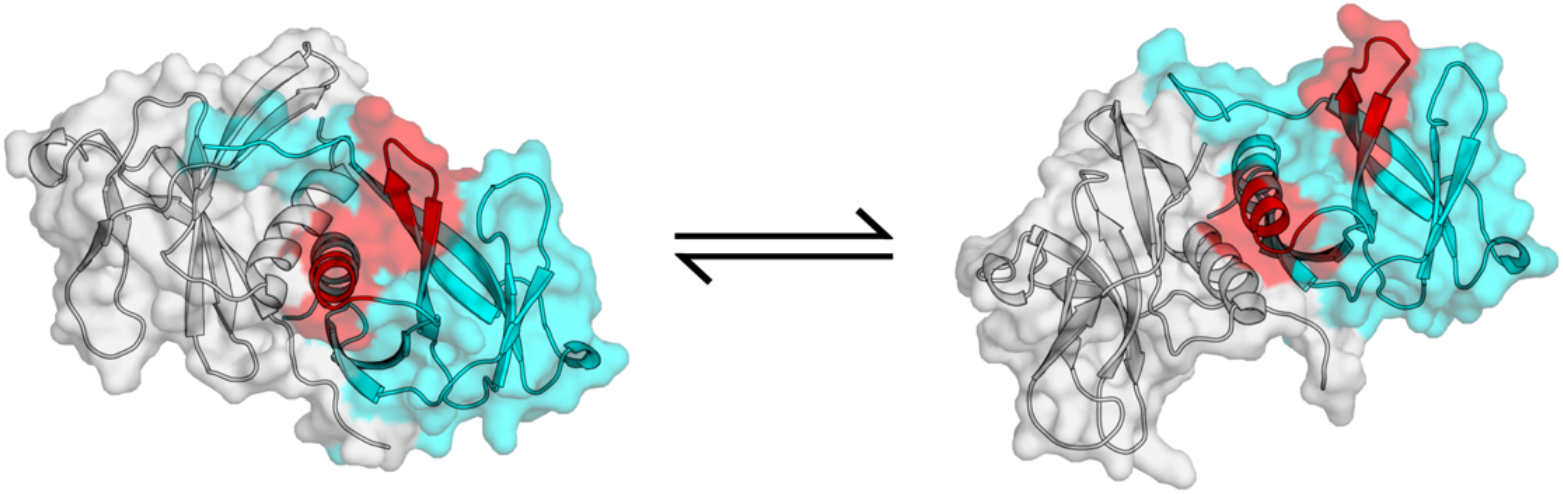
NSP9 (dimer) transition from closed to open state. Backbone is represented as a cartoon and sidechains are represented with a transparent surface. Each molecule in the dimer is identified with a unique color, gray or cyan. The residues that undergo a large conformational change to expose a cryptic pocket are highlighted in red.

**Figure S8:**
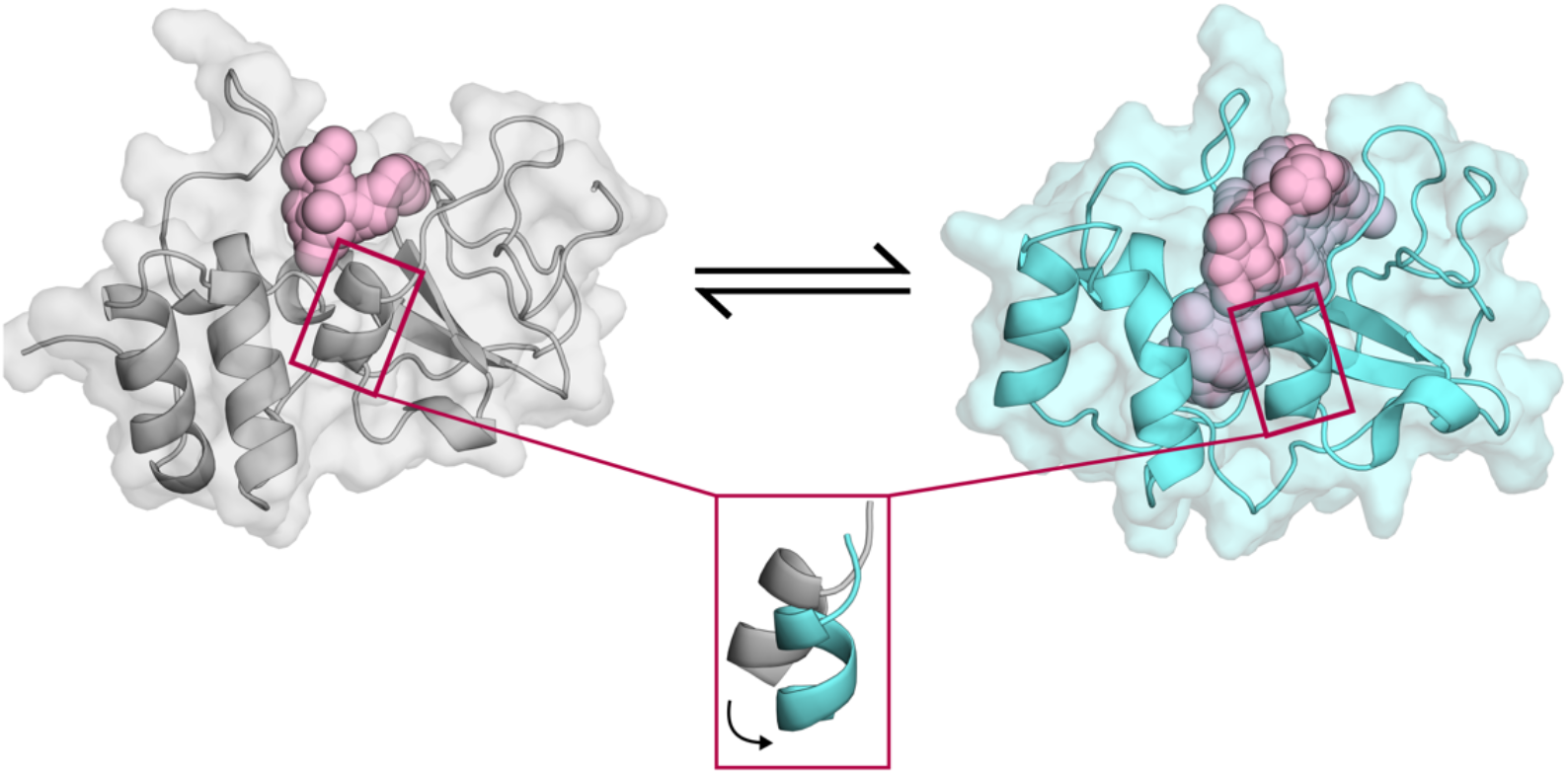
NSP10 transition from closed to open state. Backbone is represented as a cartoon and sidechains are represented with a transparent surface. Pocket volumes are highlighted with pink spheres. Here, an existing pocket is greatly expanded from the swivel of an α-helix.

**Figure S9:**
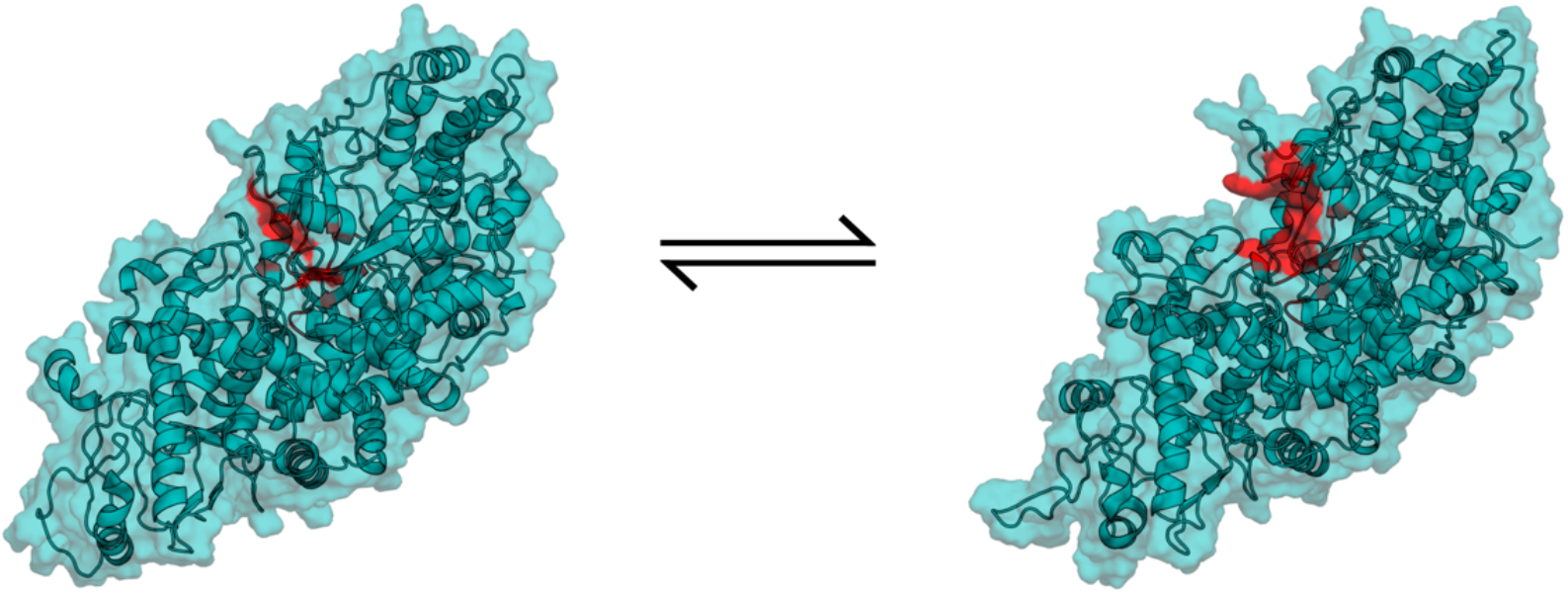
NSP12 transition from closed to open state. Backbone is represented as a cartoon and sidechains are represented with a transparent surface. The residues that undergo a large conformational change to expose a cryptic pocket are highlighted in red.

**Figure S10:**
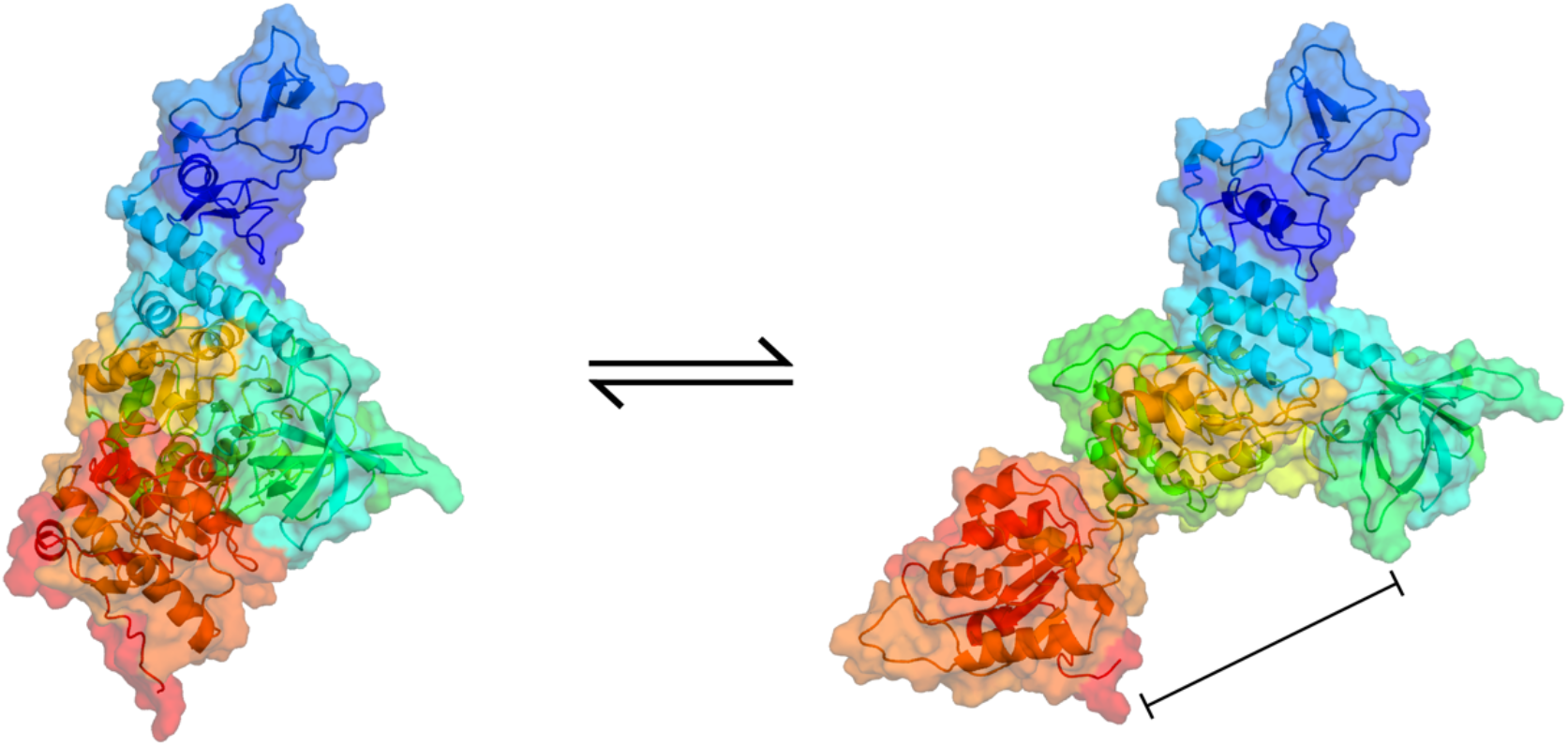
NSP13 transition from closed to open state. Backbone is represented as a cartoon and sidechains are represented with a transparent surface. The protein is colored by residue number following a rainbow and highlights the various domains. Here, we observe a large domain motion between domains 1A and 2A, which may be relevant for nucleotide binding.

**Figure S11:**
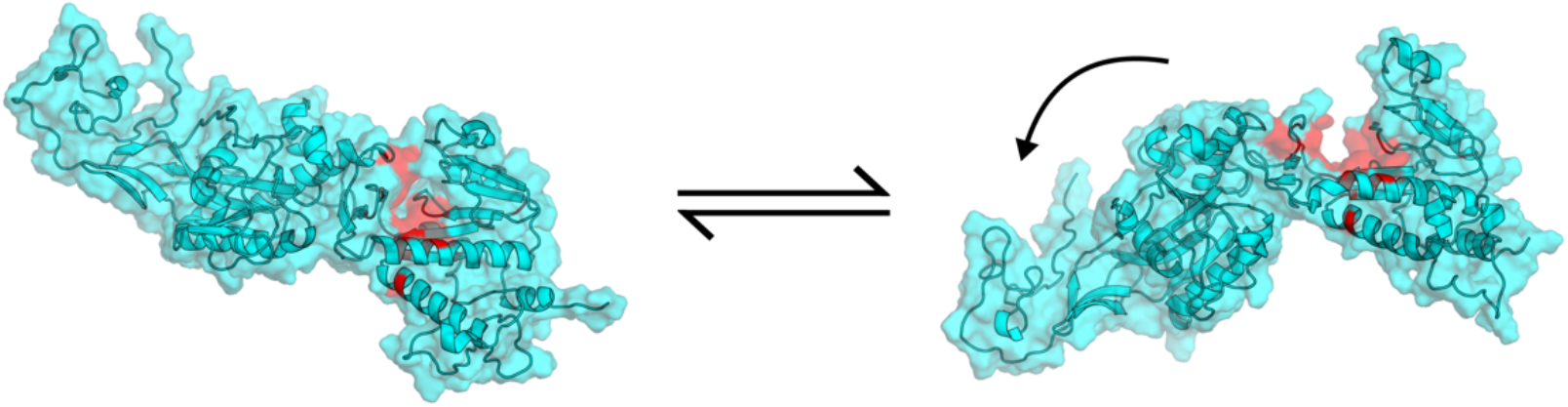
NSP14 transition from closed to open state. Backbone is represented as a cartoon and sidechains are represented with a transparent surface. The residues that undergo a large conformational change to expose a cryptic pocket are highlighted in red.

**Figure S12:**
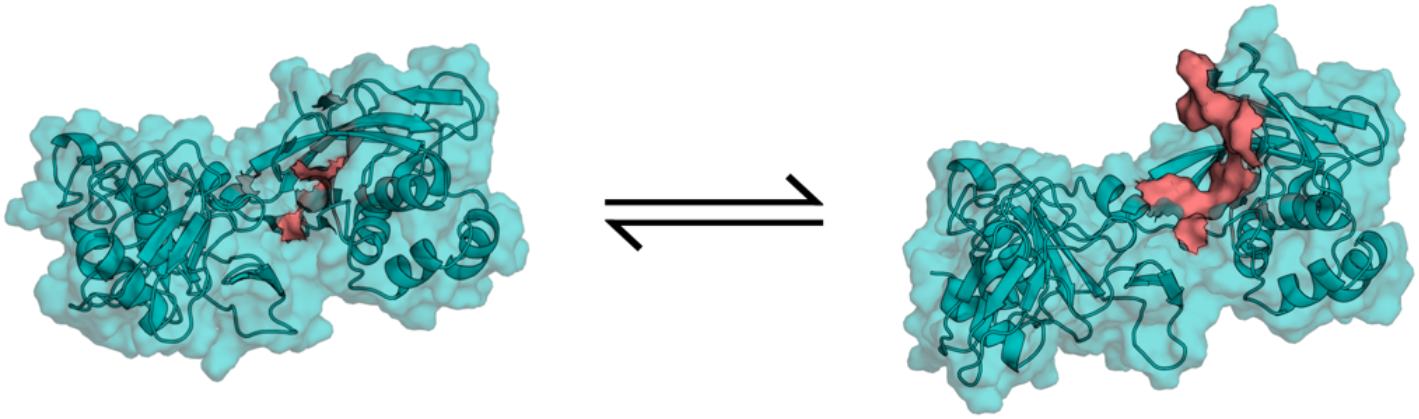
NSP15 transition from closed to open state. Backbone is represented as a cartoon and sidechains are represented with a transparent surface. The residues that undergo a large conformational change to expose a cryptic pocket are highlighted in red.

**Figure S13:**
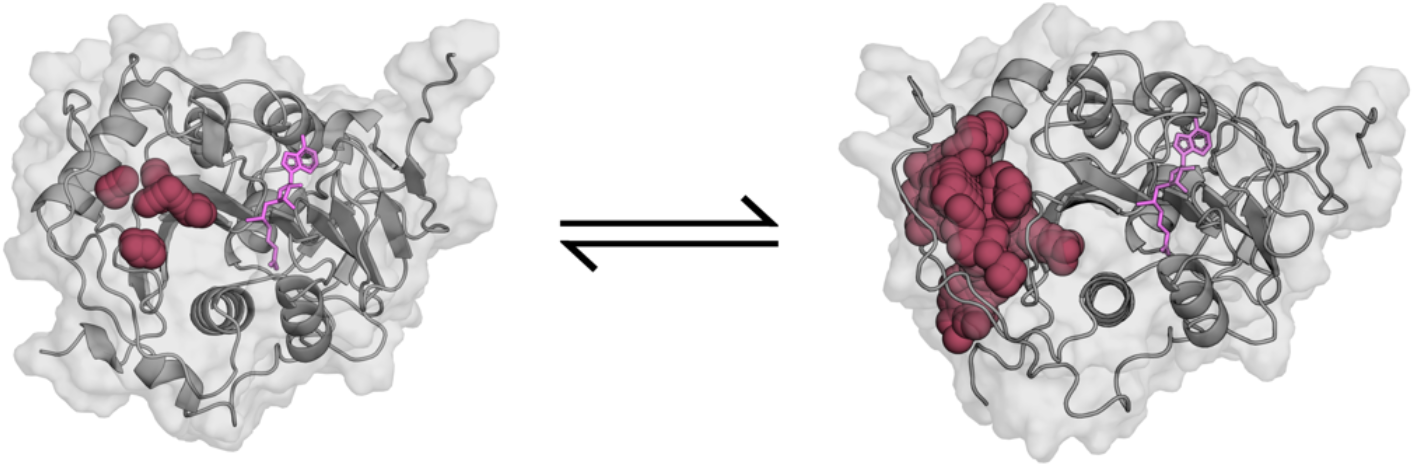
NSP16 transition from closed to open state. Backbone is represented as a cartoon and sidechains are represented with a transparent surface. Pocket volumes are highlighted with maroon spheres. SAM cofactor is shown with pink sticks.

**Figure S15:**
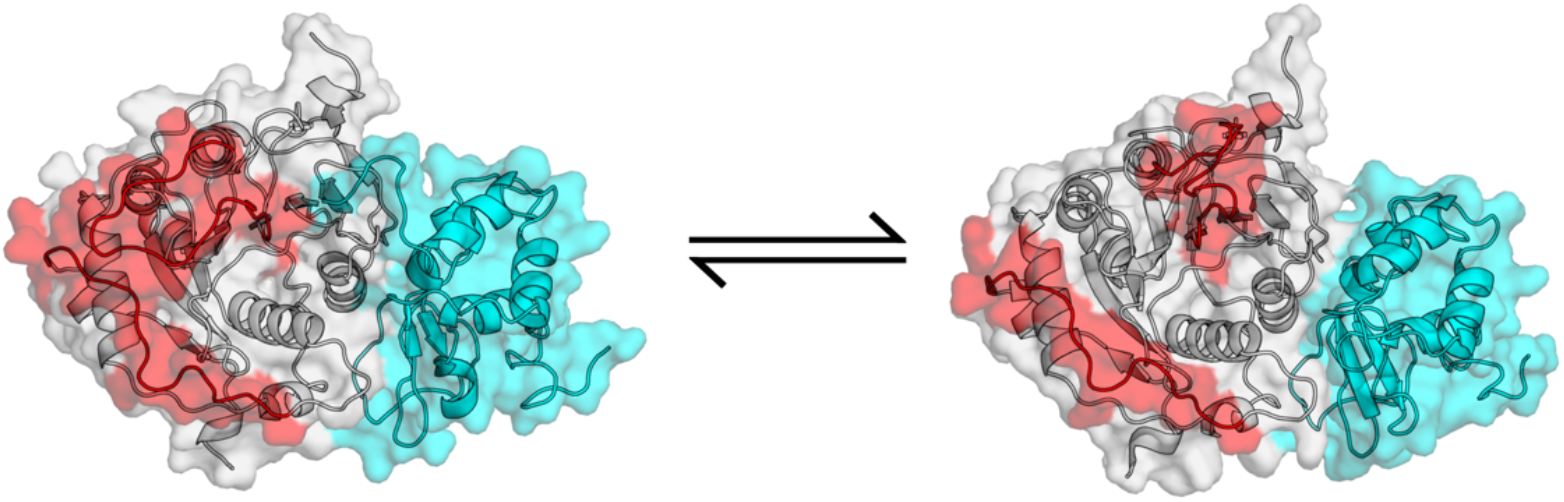
NSP10/NSP16 (complex) transition from closed to open state. Backbone is represented as a cartoon and sidechains are represented with a transparent surface. Each molecule in the complex is identified with a unique color, gray (NSP16) or cyan (NSP10). The residues that undergo a large conformational change to expose a cryptic pocket are highlighted in red.

**Figure S16:**
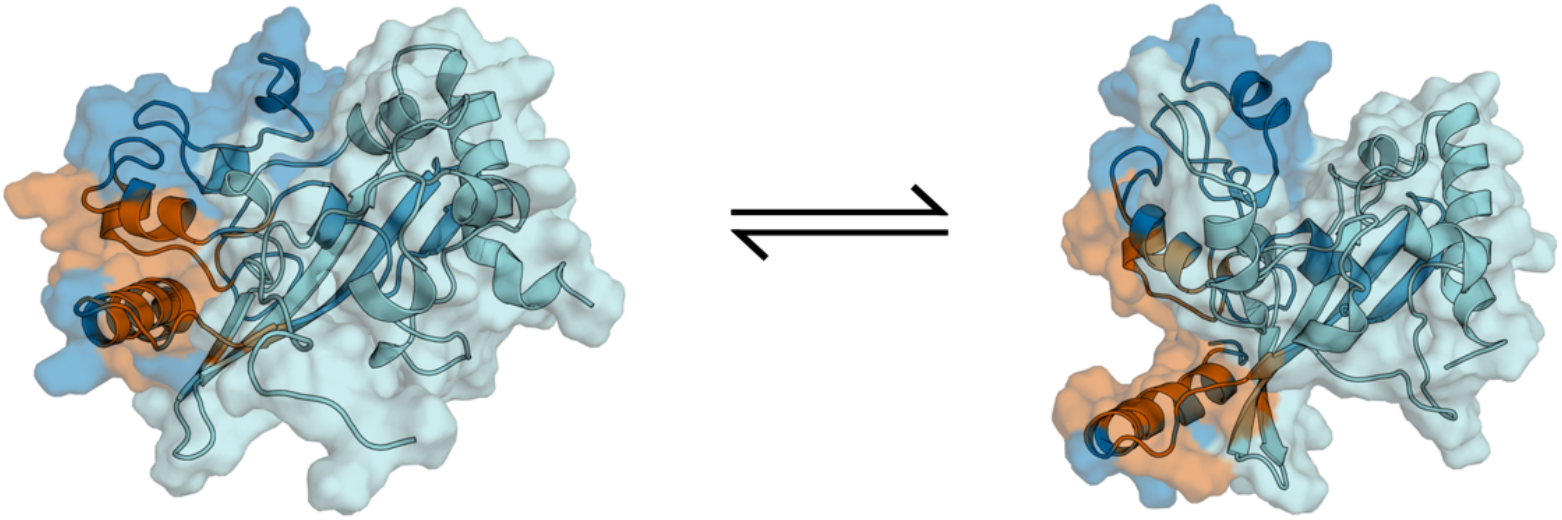
Nucleoprotein dimerization domain transition from closed to open state. Backbone is represented as a cartoon and sidechains are represented with a transparent surface. The residues that undergo a large conformational change to expose a cryptic pocket are highlighted in orange.

**Figure S17:**
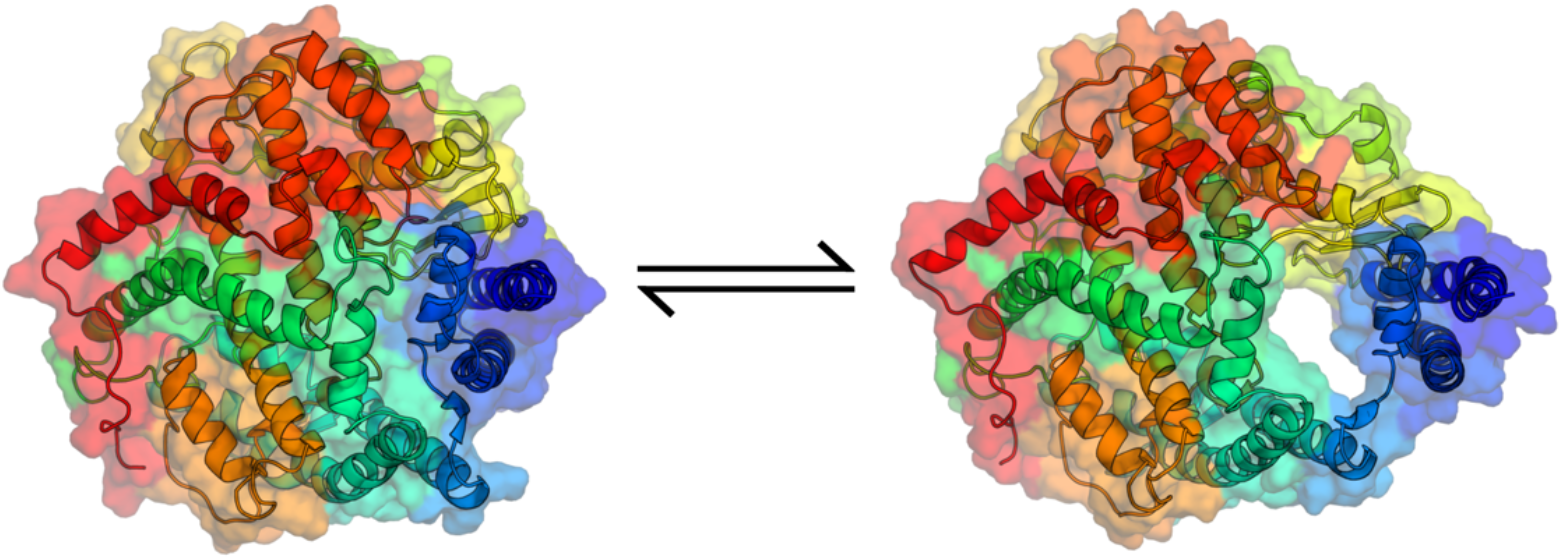
Human ACE2 transition from closed to open state. Backbone is represented as a cartoon and sidechains are represented with a transparent surface. The protein is colored by residue number following a rainbow. Pocket is proximal to the region that binds to SARS-CoV-2 spike protein.

**Figure S18:**
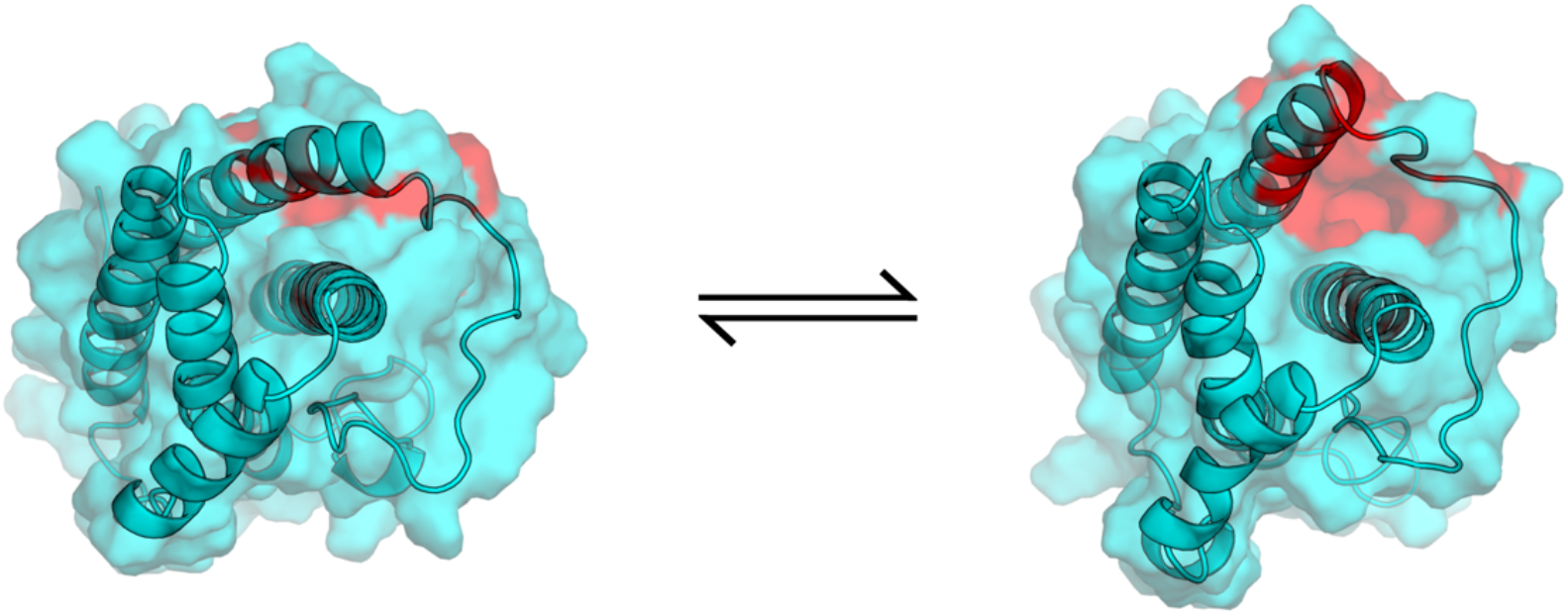
Human IL6 transition from closed to open state. Backbone is represented as a cartoon and sidechains are represented with a transparent surface. The residues that undergo a large conformational change to expose a cryptic pocket are highlighted in red.

